# Multifocal, multiphenotypic tumours arising from an MTOR mutation acquired in early embryogenesis

**DOI:** 10.1101/2023.12.12.570785

**Authors:** Clarissa N. Pacyna, Madhanagopal Anandapadamanaban, Kevin W. Loudon, Iain M. Hay, Olga Perisic, Ruoyan Li, Matthew Byrne, Laura Allen, Kirsty Roberts, Yvette Hooks, Anne Y. Warren, Grant D. Stewart, Menna R. Clatworthy, Sarah A. Teichmann, Sam Behjati, Peter J. Campbell, Roger L. Williams, Thomas J. Mitchell

**Author notes:** These authors contributed equally to this work (Clarissa N. Pacyna, Madhanagopal Anandapadamanaban). These corresponding authors contributed equally to this work (Roger L. Williams, Thomas J. Mitchell).

## Abstract

Embryogenesis is a vulnerable time. Mutations in developmental cells can result in the wide dissemination of cells predisposed to disease within mature organs. We characterised the evolutionary history of four synchronous renal tumours from a 14-year-old girl, timing their shared origin to a multipotent embryonic cell committed to the right kidney, around 4 weeks post-conception. Their shared *MTOR* mutation, absent from normal tissues, enhances protein flexibility, which enables a FAT domain hinge to dramatically increase activity of mTORC1 and mTORC2. Developmental mutations, not usually detected in traditional genetic screening, have vital clinical importance in guiding prognosis, targeted treatment, and family screening decisions for paediatric tumours.

## Main text

Four renal tumours were identified incidentally in the right kidney of a 14-year-old girl (Fig. 1a); the left kidney was radiologically normal. She underwent a radical nephrectomy and has remained free of disease as of writing. Two of the lesions (A and D) had the typical histological appearances of chromophobe renal cell carcinomas (chRCC) and two (C and E) were renal oncocytomas (RO) (Fig. 1b). Both of these tumour types, thought to be derived from collecting duct type A cells, are exceedingly rare in young patients, even with the presence of a predisposing germline mutation in *FLCN* (Birt-Hogg-Dubé syndrome).^1^ This is just the eighth case of a child with a RO, one of about twenty reported paediatric chRCC cases, and the first report of a young patient with both lesions occurring in the same kidney^2–4^. Standard clinical genetic testing detected no causative germline mutations.

**Fig. 1:**
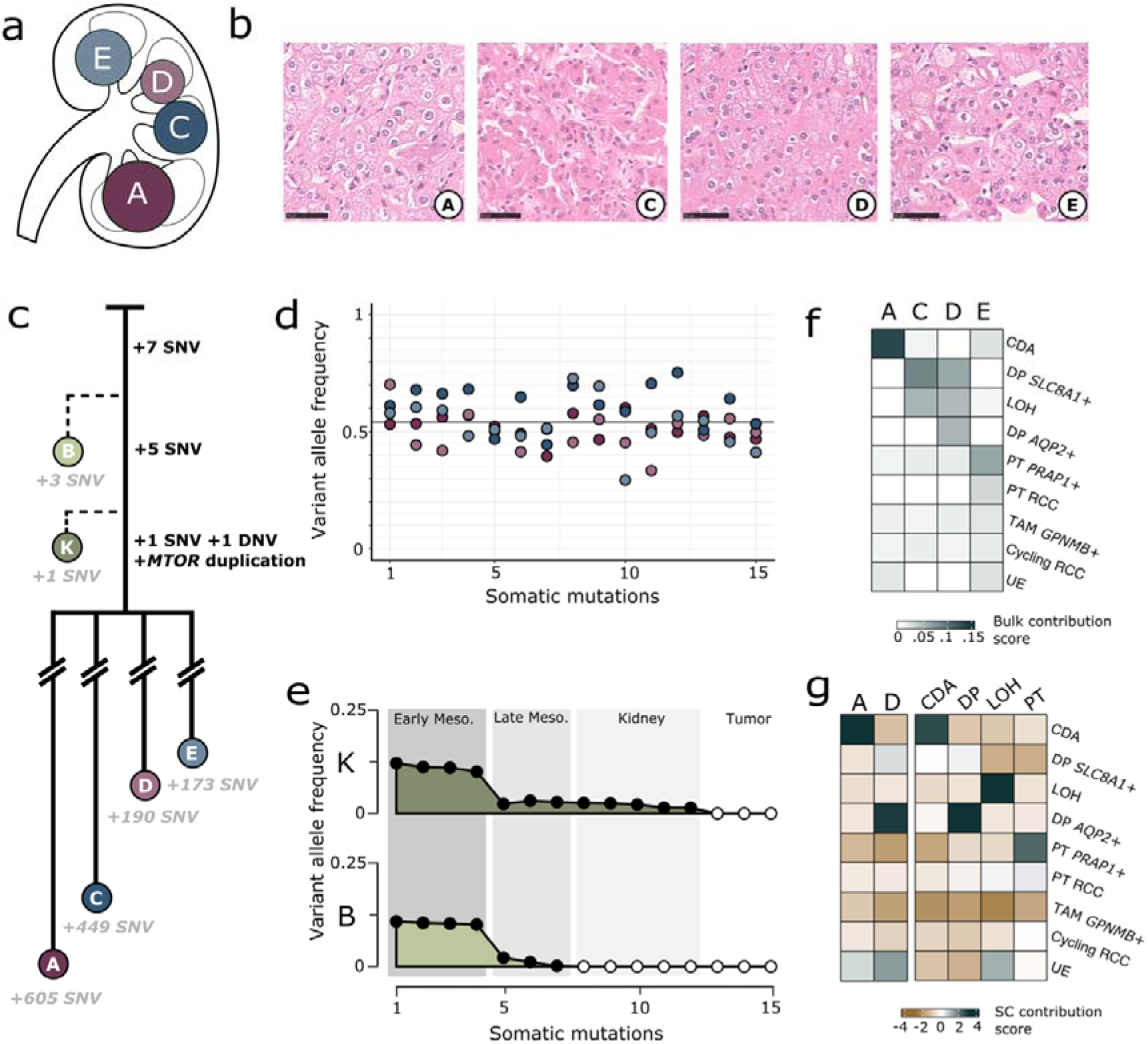
Phylogenetic and transcriptomic analysis of the patient’s four renal tumours. **(a)** The four tumours are dispersed throughout the kidney. **(b)** Tumour A is a standard RO. Tumour C is a chRCC with cellular extensions into the kidney and its neighbouring tumour. Tumour D is a RO with some nuclear atypia and foamy macrophages. Tumour E is a chRCC with cellular extensions into the kidney. Scale is 50 μm. **(c)** Blood and normal kidney (B and K respectively) have the expected number of shared and independent mutations. Once the four tumours acquired the *MTOR* duplication, their lineages split and cancer progenitors seeded throughout the kidney and evolved separately. **(d)** The fifteen shared SNVs are clonal in the four tumours with a displayed mean VAF of 0.54. **(e)** The 15 shared clonal tumour mutations in normal kidney (K) and blood (B) have VAFs that begin at about 0.125 and decrease to zero, reflecting the stage in development in which they were likely acquired. **(f)** Deconvolution of bulk RNA-seq data reveal that each tumour’s transcriptome best resembles that a different renal cell type. The top scored renal cell types (CDA: collecting duct type A, DP *SLC8A1*+: distal principal *SLC8A* positive, LOH: loop of Henle, PT *PRAP1*+: proximal tubule *PRAP1* positive), two tumour cell types (PT RCC: proximal tubule RCC, cycling RCC: cycling RCC), and immune cell type (TAM *GPNMB+*: tumour associated macrophage *GPNMB* positive) are reported for each bulk tumour sample. **(g**) Logistic regression analysis of single cell RNA-seq clusters for tumour A and D most closely resemble collecting duct type A and distal principal cells, respectively. In the second panel, normal single cell clusters are defined from similarity to collecting duct type A (CDA), distal principal (DP), Loop of Henle (LOH), and proximal tubule (PT) cell types. Row labels same as in **f**.

We evaluated the phylogenetic relationships of these four renal tumours using whole genome sequencing. The tumours shared 13 single nucleotide variants (SNV), one double nucleotide variant (DNV) and one in-frame 12 base-pair (bp) duplication in *MTOR* (Fig. 1c). Beyond this small number of shared variants, each separate tumour had independently acquired somatic mutations with distinct copy number profiles (Extended Data Fig. 1a). The 15 shared mutations were clonal in all tumours (Fig. 1d). High-coverage whole genome sequencing (550x) of histologically normal kidney and blood showed that 7 of the 15 shared variants were detectable in both normal kidney and blood, with monotonically decreasing variant allele frequency (VAF). A further 5 variants were detectable in normal kidney but not blood, and 3 variants were not detectable in either (Fig. 1e).

We also analysed bulk and single cell transcriptomes from the four tumours. Strikingly, each tumour had a unique transcriptome resembling a different renal cell type, inferred using deconvolution from a single-cell RNA-sequencing (scRNA-seq) reference dataset from normal kidney (Extended Data Fig. 1b-c). These cell types were collecting duct type A, distal principal, loop of Henle, and proximal tubule cells (Fig. 1f). Using the two RO tumours’ unique patterns of whole chromosome loss, we also identified cells from tumours A and D in the scRNA-seq dataset. These cells’ transcriptomes recapitulated the bulk RNA-seq findings, with tumour A cells again resembling collecting duct type A and tumour D resembling distal principal cells (Fig. 1g).

Taken together, these data demonstrate that the most recent common ancestor (MRCA) of the four tumours was a multipotent embryonic cell that likely existed during the early development of the right kidney. A burden of 15 mutations would time the MRCA to well before 8 weeks post-conception (when mesodermal cells have an average of 25 mutations each^5^), to around the fourth week of embryogenesis. This is supported by the 7 mutations shared with blood, which splits from kidney when gastrulation establishes intermediate and lateral mesoderm structures in the third week.^6^ As the nephrogenic cords split into left and right pronephros between the third and fourth week of embryogenesis, it is likely that the initially affected cell was already fated to the right kidney, generating an organ-restricted cancer field effect akin to the 1953 theorisation of “synchronous” cancer incidence.^7^ Furthermore, the fact that each of the four tumours had an expression signature of a different cell type argues that the most recent common ancestor was multipotent, capable of generating several distinct functional zones of the nephron.

Of the 15 shared mutations in the most recent common ancestor, the only coding variant was an in-frame duplication of 12bp in *MTOR*. Mammalian target of rapamycin (mTOR) is a kinase that regulates cell growth and proliferation via protein mTOR Complex 1 (mTORC1) and mTOR Complex 2 (mTORC2). The mTOR pathways are commonly dysregulated in cancer, but activating mutations in *MTOR* are rarely observed outside of renal cell carcinomas.^8^ This patient’s in-frame duplication of 1455-EWED-1458 is in the helical autoinhibitory FAT domain of mTOR, in close vicinity to a slightly activating single amino acid mutation A1459P seen previously in other RCCs (Fig. 2a).^9,10^ All four tumours express the mutant *MTOR* transcript, and an unbiased gene set enrichment test of the tumour transcriptomes revealed the mTORC1 hallmark signalling pathway to be significantly upregulated (Extended Data Fig. 1d). As this exact mutation had not been previously described, we probed the functional and structural consequences of the protein variant.

**Fig. 2:**
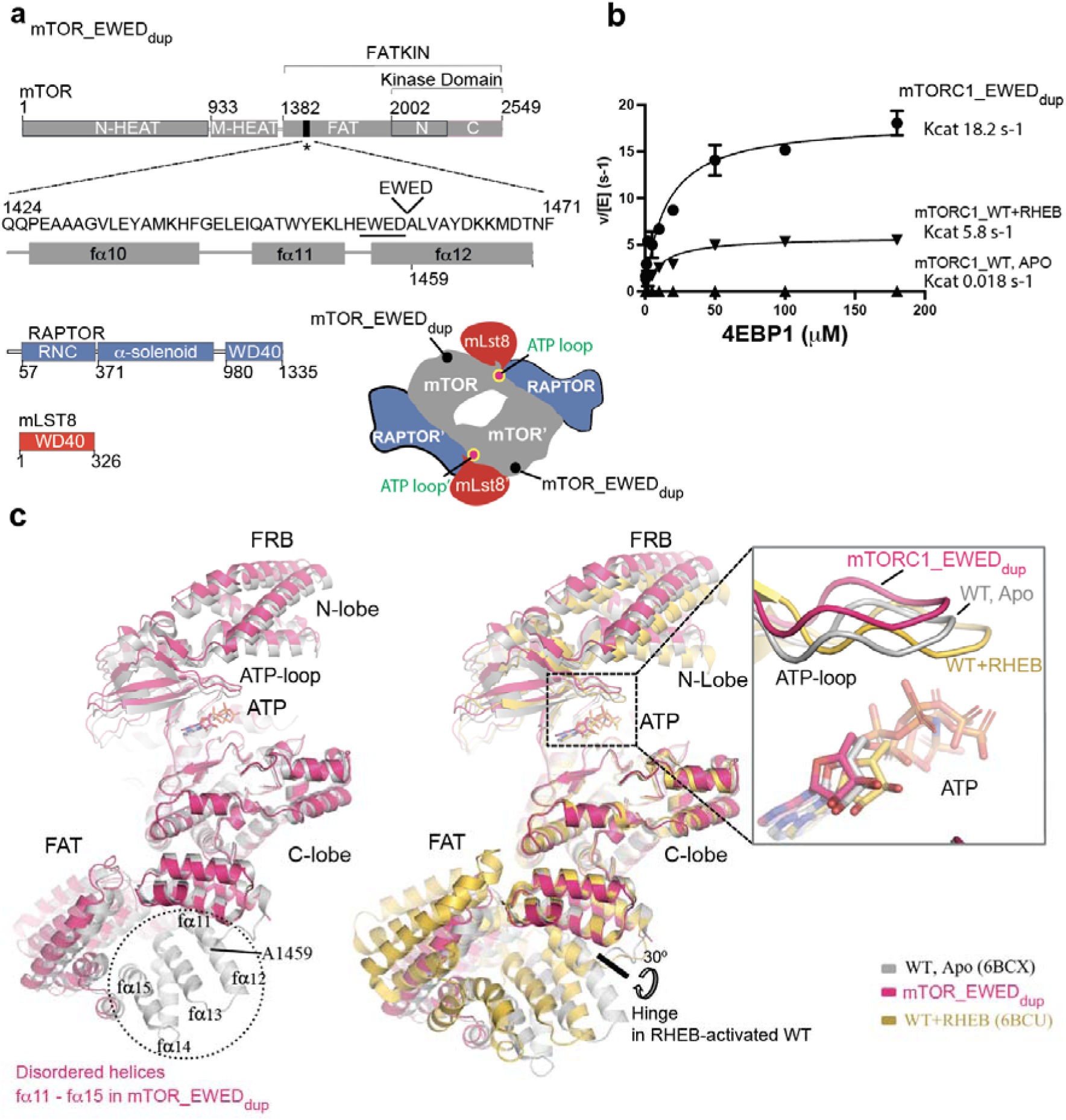
Cryo-EM structure and kinetic analysis of mTORC1_EWED_dup_ mutant. **(a)** Schematic representation of mTORC1 subunits: mTOR, RAPTOR and mLST8. The four-residue cancer-associated 1455-EWED-1458 duplication in the FAT region of mTOR is highlighted. Also shown is a schematic sketch of the subunit arrangement within a dimeric mTORC1 complex, with the regions of interest highlighted. **(b)** Kinetic analysis of 4EBP1 phosphorylation by mTORC1_EWED_dup_ mutant or by mTORC1_WT in the absence and presence of RHEB (250 µM). Graphs show means (with markers as indicated) and *k*_cat_ values using data from three independent experiments. **(c)** Cryo-EM reconstruction of mTORC1_EWED_dup_ mutant (colored in dark red) reveals the disordered region at the duplication site, corresponding to WT mTOR FAT helices (fα11-fα15). Residue A1459 in the mTORC1_WT is highlighted. Alignment of cancer-associated mTORC1_EWED_dup_ structure with mTORC1_WT apo (6BCX, gray) or bound to RHEB (6BCU, orange). The alignment was done on the C-lobe of mTOR kinase domain. Although hyperactive, the mTORC1_EWED_dup_ mutant does not undergo global conformational changes or ATP-loop realignment (shown in the inset) as observed with RHEB-activated mTORC1_WT.

First, we expressed and purified recombinant mTORC1 with the cancer-associated 1455-EWED-1458 duplication, referred to from here on as mTORC1_EWED_dup_ (Fig. 2a and Extended Data Fig. 2a,b). The thermal unfolding profile for this mutant is similar to wild type (WT), suggesting no major global destabilising events from the duplication (Extended Data Fig. 2c). Kinetic analysis showed the variant is hyperactive with a catalytic turnover (*k*_cat_) of ∼18 s^-1^, which is a great increase relative to the mTORC1_WT (*k*_cat_ of ∼0.02 s^-1^) (Fig. 2b). The activity of mTORC1_EWED_dup_ is similar to mTORC1_WT when activated by the physiological activator, RHEB (*k*_cat_ of ∼6 s^-1^), and the activity of the variant was not further increased by the addition of RHEB (Fig. 2b). Though the role of mTORC2 in tumorigenesis is less understood, we also analysed effects of the mTOR_EWED_dup_ variant on mTORC2 activity, revealing that the variant caused significant hyperactivation of this complex as well (Extended Data Fig. 3).

The cryoEM structure of the mTORC1_EWED_dup_ variant revealed a disordering of the FAT domain between residues 1442-1512, implying greater flexibility around the duplication site (Fig. 2c and Extended Data Fig. 4). Despite its hyperactivity, the mutant’s global structure resembled the apo-conformation of mTORC1_WT (apo meaning not bound to activated RHEB). This flexibility suggests that the mutant can more readily adopt a transient activated conformation, even though the equilibrium conformation resembles apo-mTORC1 (Fig. 2c and Extended Data Fig. 4). In contrast to the characteristic 30° rotation around a hinge at residue 1443 as seen in the RHEB-activated mTORC1_WT complex, the duplication variant does not elicit such a hinge motion^11^ (Fig. 2c).

Prior analysis by Xu and colleagues suggested that most *MTOR* mutations associated with renal tumours occurred in three clusters of the FAT domain (F1, F2 and F3) and in three clusters in the kinase domain.^12^ These point mutations each increased the activity of mTORC1. Furthermore, combining two mutations, one in the FAT domain and one in the kinase domain, synergized to hyperactivate mTORC1, even in the absence of RHEB.^12^ Notably, this patient’s *MTOR*_EWED_dup_ variant, present in the F1 FAT domain cluster, also hyperactivated mTORC1 to achieve a RHEB-like maximum rate (Fig. 2b), without requiring mutation in a second cluster. As this F1 cluster lies near a previously described hinge for RHEB-activated mTORC1, it is plausible that the significant disorder induced by the duplication event directly relieves auto-inhibition exerted by the FAT domain on the kinase in the basal state. The hyperactivity of the variant within mTORC2 is also consistent with the mTORC2-activating effect reported for a nearby point mutation at 1483.^13^

Together, these genomic, biochemical, and structural analyses affirm that this *MTOR* mutation, acquired early in development, is the initial tumorigenic driver. *MTOR* mutations are relatively uncommon in kidney cancer, with an incidence of 3% in chRCC and no previously reported incidence in RO tumours.^14,15^ However, the mTORC pathway has been implicated in both tumour types. Germline loss-of-function mutations in mTOR pathway gene *FLCN* in Birt-Hogg-Dubé syndrome predispose not only to chRCC and RO, but also to more aggressive clear cell and papillary RCCs.^16^ Further, recent changes to the World Health Organization (WHO) guidelines have defined a class of hybrid RO-chRCC tumours defined as “other oncocytic/chromophobe RCC” for which *MTOR* or *TSC1/2* mutations are frequently the primary driver, with suggestion that rapalogs may be an effective targeted treatment as in adult RCC patients with advanced disease.^17–19^ It is notable that the WHO guidelines highlight how *MTOR* and mTOR pathway mutations can generate a wide variety of histologically diverse tumours as we have observed in this one individual.

The standard categorisation of tumours considers germline and sporadic aetiologies as two entirely separate, opposing entities. With advances in understanding the developmental genome and with the accessibility of whole genome sequencing, there is a growing appreciation for a subset of tumours generated from some point between. As such, it may be helpful to consider tumour aetiology as a continuum from early germline to late sporadic tumours, with particular vulnerabilities through development. A number of cases of both blood cancer^20–22^ and kidney cancer^23–25^ have been described in which the first driver mutation was acquired *in utero*. What makes our case so interesting is that the distinct expression profiles across the tumours, the small number of shared mutations and the restriction to a single kidney enables us to pinpoint the most recent common ancestor to a very specific stage of development, likely a multipotent kidney progenitor cell at about 4 weeks post-conception. The timing and tolerability of a developmental driver will determine the extent of the resultant cancer field effect and the type of cancers that would eventually emerge. Because these aspects can inform prognosis and treatment, it is vital that mosaic aetiologies are further characterised along the spectrum from germline to sporadic tumours.

## Methods

### Samples and sequencing

Human material was obtained from patients enrolled in an UK NHS REC approved study (reference 16/EE/0394). After radical nephrectomy, four tumour and one normal kidney samples were biopsied and divided into fresh tissue for dissociation and fresh frozen for subsequent WGS and bulk RNA-seq. An additional perioperative blood sample was taken. DNA, RNA, and single cell suspension were extracted from the resulting fresh frozen samples.

DNA was extracted from DNA libraries of 150 bp length were prepared for Illumina NovaSeq 6000 paired end sequencing with a 500 bp insert size. The four tumour samples were sequenced to an average coverage of 76.73x, and the normal kidney and blood samples were initially sequenced to similar depth, then topped up to 496.69x and 531.69x, respectively, to identify shared variants.

Bulk RNA libraries for the five renal samples were prepared with 450-500 bp fragment size for 150 bp paired end sequencing with Illumina 2500 HiSeq chemistry to a mean coverage of 68.66x.

Single cells were dissociated in PBS, loaded, barcoded, and processed for 10X library preparation using the same protocol as reported in Li, et al*. Cancer Cell* 2022.^26^ Samples were sequenced using Illumina NovaSeq 6000 paired end machines.

### Histology preparation and processing

Paxgene fixed tissue blocks were stained with H&E and imaged with a Hamamatsu NanoZoomer S60 slide scanner.

### Genomic data processing, variant calling, and phylogeny analysis

Genomic data were aligned to the GRCh 37d5 reference genome with the Burrows-Wheeler transform (BWA-MEM). Chromosomal copy number changes were identified using both ASCAT NGS (https://github.com/cancerit/ascatNgs/tree/dev) version 4.3.3 and Battenberg (https://github.com/cancerit/cgpBattenberg) version 3.5.3 (Supplemental Table 3 and Extended Data Fig. 1a). SNVs were called using CaVEMan (Cancer Variants through Expectation Maximisation, https://github.com/cancerit/CaVEMan) version 1.15.1 and indels were called using an in-house Pindel build (https://github.com/cancerit/cgpPindel) version 3.3.0, both run in unmatched mode calling variants against a simulated reference genome dataset. We selected SNVs with fewer than half of supporting reads as clipped (CLPM = 0) and a median alignment score greater than or equal to 140 (ASMD ≥ 140). Mapping quality and base quality cut-offs were set to minimums of 30 and 25, respectively.

We identified germline and somatic variants using previously described exact binomial testing.^23^ A Shearwater-like filter was deployed to find probable sequencing artefacts by comparing called SNVs to an internal panel of 21 in-house, unrelated renal WGS samples. To select variants shared by all four tumours, we performed an additional Fisher’s exact test comparing renal variants (tumour and normal) to blood, using a cut-off of *p* < 0.01 to select for somatic, non-artefactual SNVs enriched in the tumours.

The phylogeny was drawn using both a manual and a Dirichlet process with a 10,000 burn-in on the filtered somatic SNVs.

In generating Fig. 1d, we assumed shared variants to be clonal and accordingly adjusted variant allele frequencies (VAF) to a theoretical mean of 0.5 while accounting for tumour-specific chromosome copy number changes. Raw and adjusted VAFs are included in Supplementary Table 1 and 2, respectively.

### Bulk transcriptomic data processing

Bulk transcriptomic data were mapped to the hg37d5 reference genome using the Ensembl 75 transcriptome with aligner STAR (https://github.com/alexdobin/STAR) version 2.5.0.

Differential expression analysis reported in Supplementary Table 4 was computed using DESeq2 version 1.30.1. We compared all four tumours against a panel of the one matched normal kidney and 10 GTEx normal kidney cortex samples from female donors aged 20-59. The test also included a batch correction between our in-house and GTEx sequencing. Patterns of gene programme and pathway changes were calculated from genes ranked by log-fold change in the GTEx panel DE analysis using fgsea (https://github.com/ctlab/fgsea) version 1.16.0 with Hallmark pathways, with all results in Supplementary Table 5 and the significantly enriched gene programmes with *padj* < 0.05 and Net Effect Size (NES) > 1.5 plotted in Extended Data Fig. 1d.

Expression of the mutated *MTOR* allele was determined using a samtools search of RNA-seq BAM files spanning the *MTOR* gene coordinates.

### Single cell transcriptomic data processing

scRNA-seq datasets were mapped with CellRanger and processed using Seurat version 4.0.4. Because oncocytoma and chromophobe tumours are known to have abnormally high numbers of mitochondria, we did not filter out cells with high mitochondrial reads and instead removed cells with unexpectedly too few (<200) or too many (>2000) expressed genes.

These filtered data were then processed using NormalizeData(), FindVariableFeatures() with VST selection method, and ScaleData() Seurat functions. To retain as many tumour cells, which may divide at a higher rate than normal cells, cycling cells were identified using the CellCycleSorting() function but not removed. RunPCA() was performed before inspecting the data. Lastly, neighbourhoods were defined using the FindNeighbors() function with 30 dimensions and cells were clustered using the FindClusters() function with a resolution of 2.1. Clusters were visualised in UMAP space (Extended Data Fig. 1b).

We then used AlleleIntegrator (https://github.com/constantAmateur/alleleIntegrator) to impute tumour cell identity by calling heterozygous SNPs from tumour DNA samples and searching for patterns of chromosome losses in single cells.^27^ Cells with allelic imbalances that matched the chromosomal copy number profiles of tumours A and D called from WGS data were labelled as tumour, which enabled identification of tumour cell clusters (Extended Data Fig. 1c).

To identify expression of the mutated *MTOR* allele, we searched scRNA-seq BAM files downsampled to *MTOR* gene coordinates for the 12 bp duplication using samtools.

### Inferring cell type identity and contribution from transcriptomic data

We used two methods of integrating bulk and single cell RNA-seq data. First, we started with bulk deconvolution using Cell Signal Analysis (https://github.com/constantAmateur/cellSignalAnalysis), comparing the four tumour bulk transcriptomes to a normal renal scRNA-seq reference.^26,28^ The top nine most similar cell type identities are shown in Fig. 1f.

Next, we trained a logistic regression model on the same renal scRNA-seq reference as previously described.^29^ The most likely cell type was manually assigned to each Seurat-defined cluster. Tumour cell clusters determined by allelic pattern imbalances are displayed in Fig. 1g along with several clusters determined by this method to be renal cell types.

### Cloning of recombinant human mTOR wild-type and EWED_dup_ constructs

For the production of human mTORC1 complex, the three subunits were cloned individually into a pCAG mammalian expression vector as described before.^30,31^ Cloning of the mTOR ‘EWED’ duplication mutant was carried out using a fragment-assembly based approach: two fragments were PCR amplified with overlaps containing the desired 12 bp _1455_EWED_1458_ duplication region at one end and overlaps with the digested vector at the other, and then they were assembled with the digested vector using NEBuilder HiFi DNA Assembly. mTOR subunit has an N-terminal tandem Strep-tag II followed by a TEV cleavage site, while RAPTOR and mLST8 subunits are without tags. Human RICTOR was PCR-amplified from IMAGE:9021161 clone, and then cloned with an N-terminal 3X Flag-tag in the pCAG vector for expression in mammalian cells. Human Sin1.1 was PCR-amplified from Addgene 73388 plasmid (gift from Taekjip Ha^32^) and then cloned into pcDNA4TO. Subsequently, the promoter-gene (SIN1.1)-terminator cassette was PCR amplified from this plasmid and cloned into a pCAG vector already containing mLST8 gene, to allow co-expression of the two proteins from a single plasmid

### Recombinant protein expression and purification

mTORC1 and mTORC2 complexes, containing either mTOR_WT or mTOR_EWED_dup_ mutant, were expressed by transient transfection of Expi293F cells grown in Expi293 media (Thermo Fisher A1435102) in a Multitron Pro shaker set at 37 °C, 8% CO2 and 125 rpm. A total of 1.1 mg DNA/L cells was co-transfected into cells at a density of 2.5 x10^6^ cells mL^-1^ using PEI (Polyethyleneimine "MAX", MW 40,000, Polysciences, 24765, total 3 mg PEI/L cells). After 52 h (mTORC1) or 68 h (mTORC2), cells were harvested by centrifugation and cell pellets were frozen in liquid N_2_.

mTORC1 was purified from cell pellets from 2 L Expi293F culture as described before,^31^ by affinity purification on a Strep-Trap HP resin, followed by Strep-tag cleavage by TEV protease overnight on the column. The cleaved protein was further purified by anion-exchange chromatography (AEX) on a 5 mL HiTrap Q column (Cytiva), concentrated with Amicon Ultra-4 100 kDa concentrators, flash frozen in liquid N2 and stored at -80 °C.

mTORC2 was purified from 2L Expi 293F cell pellets. Cells were lysed in lysis buffer consisting of 50 mM BICINE, pH 8.5, 300 mM NaCl, 2 mM MgCl_2_, 0.5 mM TCEP and the clarified cell lysate loaded onto 2 tandem Strep-Trap HP 5 ml columns. The loaded column was washed with 20 CV lysis buffer followed by 20 CV wash buffer (50 mM BICINE, pH 8.5, 150 mM NaCl, 2 mM EDTA, 0.5 mM TCEP) and protein eluted with wash buffer supplemented with 5 mM desthiobiotin. mTORC2 containing fractions were pooled, diluted to <100 mM NaCl and further purified on a 5 ml HiTrap Q column equilibrated in Q-buffer containing 50 mM BICINE, pH 8.5, 50 mM NaCl, 0.5 mM TCEP. Protein was eluted by a linear gradient of Q-buffer containing 1 M NaCl. mTORC2 containing fractions were pooled, concentrated and stored as described for mTORC1.

Human 4EBP1 was expressed as a GST-4EBP1 fusion in *E. coli* strain C41(DE3) and purified by affinity chromatography on Glutathione-Sepharose 4B beads, followed by GST-tag removal by incubating with TEV protease overnight. The cleaved 4EBP1 was passed through a Q column and the flow-through fractions containing 4EBP1 were concentrated and run on a Superdex 75 16/60 column equilibrated in 50 mM HEPES pH 8.0, 100 mM NaCl, and 1 mM TCEP.

Human RHEB was expressed as a GST-RHEB fusion in *E. coli* strain C41(DE3) and purified as described previously.^31^Briefly, GST-RHEB was purified on Glutathione-Sepharose 4B beads, followed by GST-tag cleavage with TEV protease overnight. To separate His_6_-tagged TEV protease from RHEB protein, sample was passed through a HisTrap FF column, followed by gel filtration on a Superdex 75 16/60 column. After removal of bound nucleotide by incubating the protein for 1 h on ice with EDTA buffer containing 20 mM HEPES pH 7, 100 mM NaCl, 20 mM EDTA and 1 mM TCEP, the buffer was exchanged to 50 mM HEPES pH 7, 100 mM NaCl, 5 mM MgCl_2_, 1 mM TCEP before concentrating the protein to 25 mg mL^-1^. The concentrated RHEB was then incubated with 1 mM GMPPNP (Jena Bioscience NU-401-50) for 60 min at 4 °C and the protein was flash frozen in liquid nitrogen and stored at -80 °C.

Human AKT1_D274A was generated by overlapping PCR mutagenesis using wild-type AKT1 (a gift from Thomas Leonard, Addgene plasmid 86561^33^) and cloned into a pAceBac1 vector with N-terminal His_10_-StrepII-(tev) tag. The recombinant protein was expressed in Sf9 insect cells and purified by affinity purification on Strep-Trap HP resin, followed by simultaneous overnight tag cleavage by TEV and dephosphorylation by λ-protein phosphatase (NEB, P0735). The cleaved, dephosphorylated protein was further purified by anion exchange chromatography on a 5 mL HiTrap Q column and AKT1 containing fractions were concentrated with an Amicon Ultra-4 30 kDa concentrator, flash frozen in liquid N2 and stored at -80 °C.

### mTORC1 activity assays

All reactions were performed in kinase buffer (KB) consisting of 25 mM HEPES, pH 7.4, 75 mM NaCl, 0.9 mM TCEP, 5% glycerol, 0.5 mg mL^-1^ BSA, at 30 °C for a duration of 30 to 45 min for the apo mTORC1_WT or for 2 to 4 min for the hyperactivated mTORC1_EWED_dup_ mutant or mTORC1_WT activated by RHEB. Reactions were set up by preincubating mTORC1 with 4EBP1 for 10 min on ice. After that, the reactions were equilibrated at 30 °C for 15 s, and kinase assays were started by the addition of 250 µM ATP and 10 mM MgCl_2_ (final concentrations). The reactions were stopped by the addition of 8 µL of 2.5X LDS sample buffer containing 4 mM ZnCl_2_ to 12 µL of sample. The samples were analysed by SuperSep Phos-tag (50 µM), 7.5% precast gels (FUJIFILM Wako Pure Chemical Corporation, 192-17381), with MOPS (no EDTA) running buffer supplemented with 5 mM sodium bisulphate. Western blots were performed using a 0.2 µm pore size nitrocellulose membrane (Invitrogen IB301002) and the iBlot dry blotting transfer system (Invitrogen). After the transfer, membranes were blocked with 5% Marvel in TBST buffer (100 mM Tris-HCl, 150 mM NaCl, 0.1 % Tween 20). Incubation with the primary antibody, anti-4EBP1 (Cell Signalling, Cat. No. 9452S), was done in 5% BSA in TBST at 4 °C overnight, using 1:1000 dilution of the antibody. Incubation with the secondary antibody (anti-Rabbit IgG, HRP-linked Antibody, Cell Signalling Cat. No.7074) was at room temperature for 1 h, using 1:5000 dilution of the secondary antibody. Detection was performed using a ChemiDoc Touch Imaging System (Bio-Rad). Kinetic parameters k_cat_ and K_M_ were calculated using Prism 9 and nonlinear regression fitting of the data assuming Michaelis-Menten kinetics.

### mTORC2 activity assays

mTORC2 activity assays were performed in kinase buffer (KB) consisting of 25 mM HEPES, pH 7.5, 100 mM NaCl, 0.5 mM TCEP and 5% glycerol. Reactions were set up by serial dilution of AKT1_D274A (10-120 μM) in KB, followed by pre-incubation with 0.2 μM mTORC2_WT or 0.01 μM mTORC2_EWED_dup_ for 5 min on ice. Samples were equilibrated to 30 °C and reactions started by addition of 1 mM ATP and 10 mM MgCl_2_. Reactions were terminated after 10 min by addition of 4X LDS sample buffer containing 8 mM ZnCl_2_. For each reaction, a sample volume equivalent to 1 μg of AKT1 was analysed by SuperSep Phos-tag gel electrophoresis as described for mTORC1. Phosphorylated AKT was visualised by Coomassie staining (InstantBlue, Abcam) and imaged using a ChemiDoc Touch Imaging System. The fraction of phosphorylated AKT was determined by densitometry and kinetic parameters calculated as for mTORC1.

### Cryo-EM sample preparation

Purified mTORC1 (1 µM), 4EBP1 (20 µM) and 1 mM AMPPNP (Jena Bioscience NU-407-10) were mixed in ∼300 µL and incubated for 1 h on ice. The sample was crosslinked with 0.2 mM BS3 for 15 min at 4 °C, followed by further crosslinking with 0.03% glutaraldehyde (Sigma G5882) (added from a 1% glutaraldehyde stock in buffer A containing 50 mM HEPES pH 7.5, 0.1 M NaCl, 1 mM TCEP, 10% glycerol), for 15 min at 4 °C. The reaction was quenched by the addition of Tris pH 8.0 (final concentration 100 mM). The sample was then immediately subjected to a gradient centrifugation on a 12 mL gradient of 10-30 % glycerol in 50 mM HEPES pH 7.5, 0.1 M NaCl, 1 mM TCEP) preformed in a SW40 rotor tube (Ultra-Clear, Beckman 344060) using a gradient maker (Biocomp Instruments). The sample was centrifuged in an SW40 rotor (Beckman) at 33,000 rpm for 16 h. After centrifugation, 0.40 mL fractions were collected, analysed by SDS-PAGE, and the fractions containing crosslinked material were pooled, and concentrated to 500 µL using an Amicon Ultra-15 100 kDa concentrator. Cross-linked mTORC1_EWED_dup_ variant was further run on a Superose 6i 10/300 column equilibrated in 50 mM HEPES pH7.5, 250 mM NaCl, 5 mM MgCl2 and 1 mM TCEP and the peak fractions were concentrated to 0.8 OD_280_ and incubated further with 1 mM AMPPNP and 20 µM of 4EBP1 for 10 min and used immediately for cryo-EM grid preparation.

### Cryo-EM data collection and processing

UltraAuFoil R 1.2/1.3 (Au 300 mesh) grids were glow-discharged using an Edwards Sputter Coater S150B for 1 min at 40 mA. A 3 μL aliquot of freshly prepared, crosslinked mTORC1_EWED_dup_ at a concentration of 0.5 mg mL^-1^ was added to the grids and blotted immediately for 3.5 s at 14 °C (95% humidity) and then plunge-frozen in liquid ethane using a Vitrobot (Thermofisher). A total of 11,197 micrographs were acquired on a FEI Titan Krios electron microscope operated at 300 keV. Zero-energy loss images were recorded on a Gatan K3 Summit direct electron detector operated in super-resolution mode with a Gatan GIF Quantum energy filter (20 eV slit width), using EPU for automated collection. Images were recorded at a magnification of 105,000 (calibrated pixel size of 0.86 Å), with a dose rate of ∼16 electrons/Å^2^/s. An exposure time of 2.3 s was fractionated into 50 movie frames, giving a total dose of 50 electrons/Å^2^. For data collection, the defocus-range was set to -2.6 to -1.2 µm.

All image-processing steps were done using the RELION 4 software package^34^, which includes Gctf,^35^ MotionCor2^36^ and ResMap^37^. A total of 11,197 micrographs were processed using GPU-accelerated MotionCor2 to correct for electron beam-induced sample motion, while the contrast transfer function (CTF) parameters were determined using Gctf. Particles were picked using RELION autopicking. In total, 1,069,382 particles were extracted with a particle box size of 512 by 512 pixels. Two rounds of reference-free 2D classification (using a mask with a diameter of 352 Å) resulted in a selection of 285,221 particles. This set of particles was subjected to a 3D classification over 25 iterations in point group C_1_, using a low-pass filtered (40 Å) ab-initio reference, which was created from the de-novo 3D model generated by the SGD algorithm in Relion4. Selection of reasonably looking classes by visualisation in Chimera and by paying attention to the rotational and translational accuracies for six classes reduced the number of particles to 264,890 sorted into four 3D classes. Without providing a mask around the mTORC1 complex, 3D auto-refinement of these particles, with C1 symmetry, led to a reconstruction of 4.2 Å resolution, based on the gold-standard FSC = 0.143 criterion.^38,39^ To correct for beam-induced particle movements, to increase the signal-to-noise ratio for all particles, and to apply radiation-damage weighting, the refined particles were further ‘polished’ using the Bayesian approach implemented in Relion 4.0. Following this step, another 3D auto-refinement using a mask around the mTORC1 complex as well as applying solvent-flattened FSCs yielded a 4.0 Å resolution reconstruction (FSC = 0.143 criterion). After a CTF- and beamtilt-refinement for the estimation of per-particle defocus and beam-tilt values for the complete set of selected particles, a 3D auto-refinement resulted in a 4.0 Å resolution reconstruction (FSC = 0.143 criterion). After correction for the detector modulation transfer function (MTF) and B-factor sharpening (sharpened with a negative B-factor as listed in Supplementary Table 6), the post-processed map was used for inspection in Chimera^40^ and model building in Coot.^41^ Local resolutions were estimated using ResMap. The 3D FSC server was used to determine the directional FSC and sphericity of the maps (Extended Data Fig. 4b)^42^. One of the mTORC1 protomer possess better EM densities compared to the other protomer. The mTORC1_EWED_dup_ dimer lacks significant densities in one of the RAPTOR molecules at the HEAT and WD40 domains. The region of FAT domain around _1455_EWED_1458_ duplication shows extreme flexibility and lacks EM densities in both mTOR_EWED protomers. Focused classification with signal subtraction of the better protomer yielded 178,379 particles. A 3D refinement of this set of particles led to a reconstruction at 3.4 Å resolution. A focused classification with signal subtraction of the better protomer particles with the masks covering the mTOR residues 769-2549 and RAPTOR residues 57-366 (mTORΔN-RAPTORΔC), and subsequent refinement of mTORΔN-RAPTORΔC yielded a reconstruction at 3.1 Å (Supplementary Table 6).

### Cryo-EM structure refinement and validation

The model for one protomer of the apo mTORC1_WT structure (6BCX) was rigid body fit into the locally refined mTORC1_EWED_dup_ EM protomer density and into the focused refined mTORΔN-RAPTORΔC density. The models were then manually adjusted using Coot. The StarMap plugin for ChimeraX^43^ was used to generate Rosetta scripts for rebuilding and refining the model.^44^ One of the half-maps was lowpass filtered and used for Rosetta refinement. The other half-map was used for FSC validation. The refined model was manually checked in COOT and B-factor refinement was performed using Rosetta. Further real space refinement was carried out in Phenix (version 1a, 4620)^45^ with default settings and the following additions: (1) a nonbonded weight of 1,000 was used; (2) rotamer outliers were fit with the target ‘fix_outliers’. MolProbity^46^ was carried out for validation and manual adjustments were made in COOT followed by re-refinement. Additional validation of the model was performed in Phenix Validation. The above refined protomer model was then rigid-body fit into both protomers of the mTORC1_EWED_dup_ dimer map, followed by the same refinement and validation procedure (Supplementary Table 6).

### Thermal stability assay

Before performing thermal stability assays, defrosted samples were gel filtered on a Superose 6 Increase (10/300) column equilibrated in 50 mM HEPES, pH 7.5, 200 mM NaCl, 1 mM TCEP. Thermal unfolding was followed by differential scanning fluorimetry (DSF) by measuring intrinsic protein fluorescence using a Prometheus NT.48 (NanoTemper Technologies). Aliquots (10 μl) of purified mTORC1_WT or mTORC1_EWED_dup_ variant were loaded into the Prometheus capillaries (Cat. PR-C002, NanoTemper Technologies), and the fluorescence intensity at 330 nm and 350 nm was recorded as a function of temperature from 10 °C to 90 °C. The experiment was repeated two times with three replicates per sample. Melting points were calculated using PR.ThermControl.

### Data and Code Availability

DNA, bulk RNA, and scRNA sequencing data are available in EGA projects EGAD00001011645, EGAD00001011646, and EGAD00001011647, respectively. Genomic analysis code, bulk RNA-seq counts, and ASCAT outputs are available at https://github.com/cpacyna/devHitchhiking. scRNA-seq counts and metadata are shared on Mendeley Data at doi:10.17632/6fg25sm5g8.1. PDB coordinates are deposited with the PDB under accession numbers 8RCH (mTORC1_EWED_dup_ dimer, overall refinement), 8RCK (mTORC1_EWED_dup_ One protomer copy) and 8RCN (mTORC1_EWED_dup_ focused refined mTORΔN-RAPTORΔC). CryoEM maps are deposited with the Electron Microscopy Databased under accession numbers EMD-19052 (mTORC1_EWED_dup_ dimer, overall refinement), EMD-19053 (mTORC1_EWED_dup_ One protomer copy) and EMD-19056 (mTORC1_EWED_dup_ focused refined mTORΔN-RAPTORΔC).

## Acknowledgements

We thank Martin Prete for help with the in-house scRNA-seq analysis server, Matthew Young for assistance with alleleIntegrator, and Manuel Medrano for comments on the manuscript draft. We thank Stephen McLaughlin (LMB Biophysics facility) for help with biophysics measurements. We thank Bilal Ahsan, Giuseppe Cannone, Grigory Sharov, Anna Yeates, and Shaoxia Chen at the MRC-Laboratory of Molecular Biology Electron Microscopy Facility for help with electron microscopy data collection. We thank Jake Grimmett, Toby Darling, and Ivan Clayson (LMB Scientific Computing) for IT support. We thank Keitaro Yamashita, Vish Chandrasekaran, and Ranagana Warshamanage for advice. We acknowledge Diamond for access and support of the cryoEM facilities at the UK national electron Bio-Imaging Center (eBIC), proposal BI23268-85.

## Extended Data Figures

**Extended Data Fig. 1:**
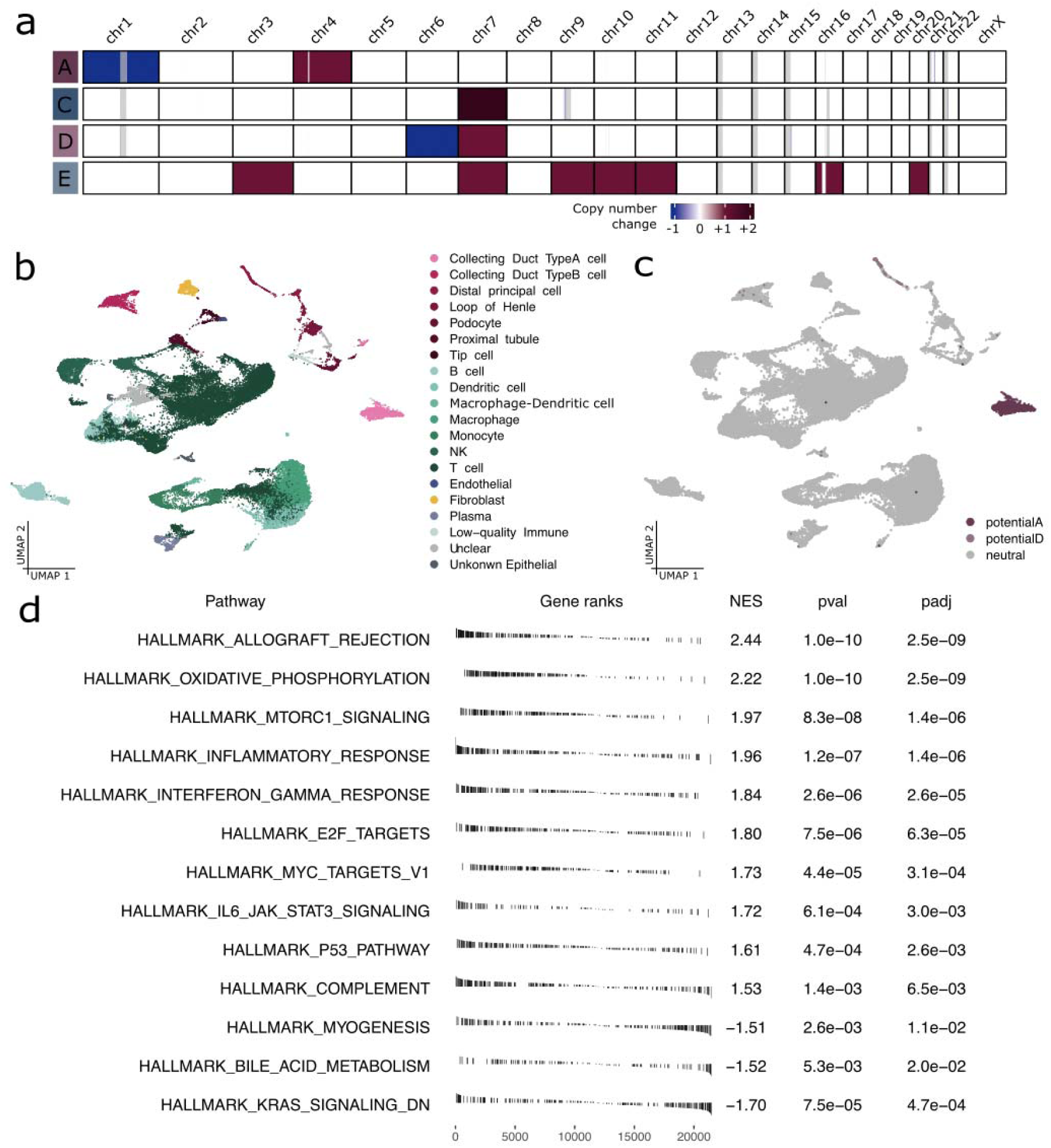
Tumour aneuploidy and transcriptomic profiles. **(a)** Each tumour has a distinct copy number change profile. **(b)** Cells are clustered and displayed in UMAP space, with each cluster labelled with a cell type identity inferred from logistic regression modelling. **(c)** Cells harbouring copy number changes that match oncocytomas A and D, highlighted in dark and light pink respectively, are grouped into two distinct clusters. **(d)** Gene set enrichment analysis comparing the four tumours to a normal GTEx panel reveals significantly up- or down-regulated Hallmark pathways, notably including mTORC1 signalling upregulated in the four tumours. Positive net effective score (NES) corresponds to upregulation in tumour.

**Extended Data Fig. 2:**
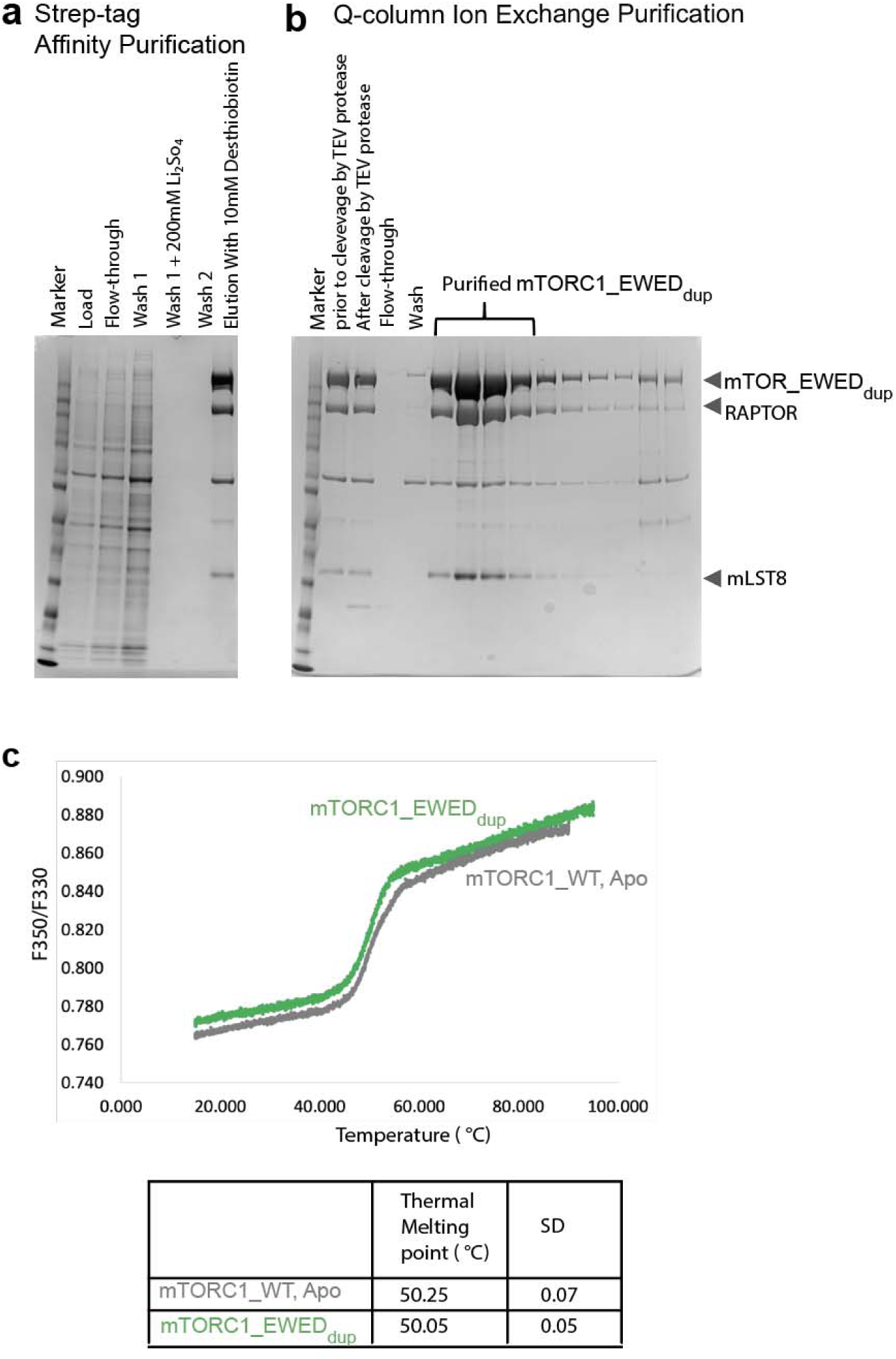
mTORC1 purification and characterization. **(a,b)** SDS-PAGE showing the purification of mTORC1_EWED_dup_ variant. First step is strep-affinity purification **(a)**, followed by Ion Exchange Chromatography **(b)**. **(c)** Thermal stability of mTORC1_WT and mTORC1_EWED_dup_ variant measured by differential scanning fluorimetry (DSF).

**Extended Data Fig. 3:**
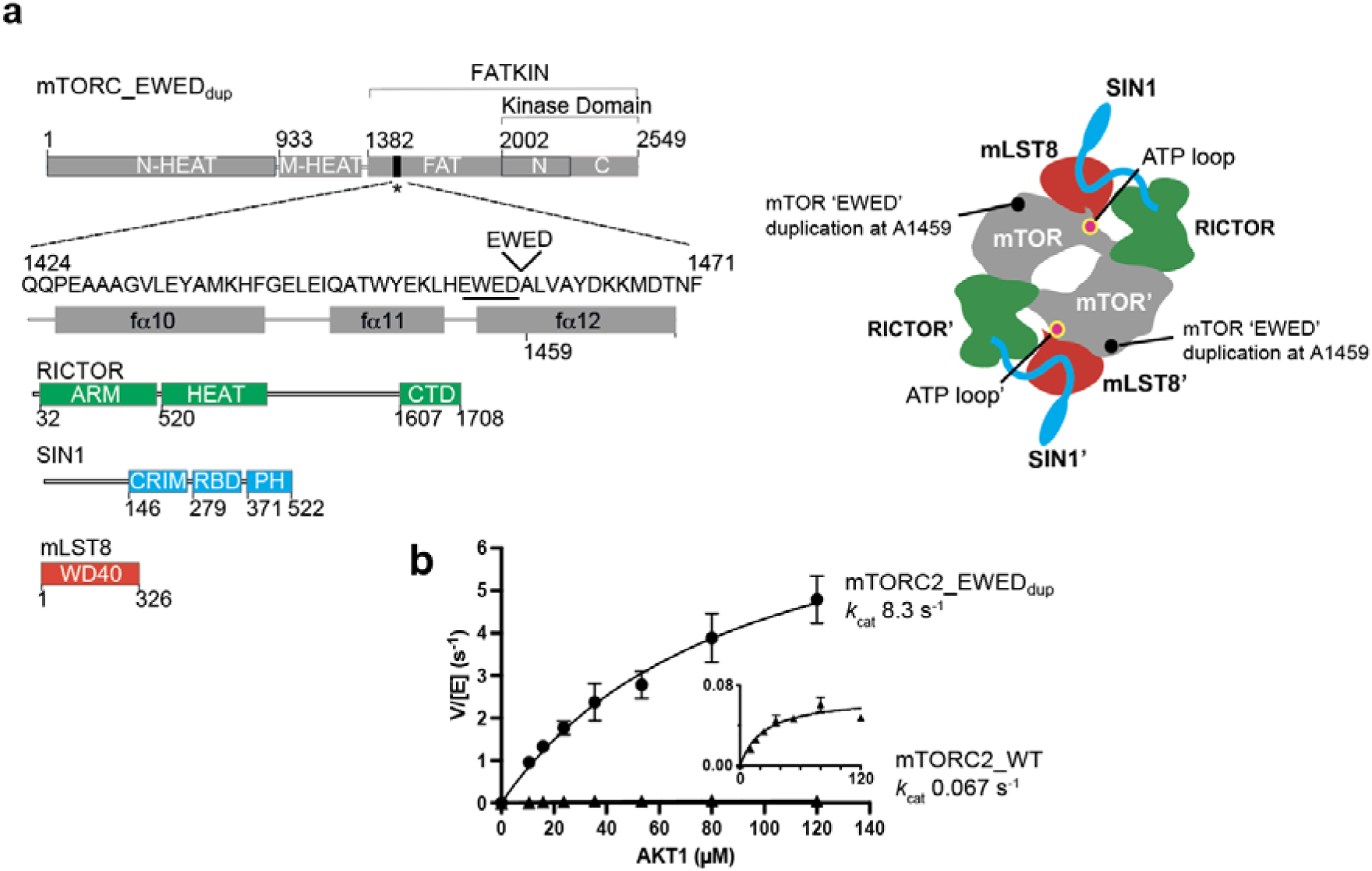
mTOR_EWED_dup_ mutation hyperactivates mTORC2. **(a)** Schematic representation of mTORC2 subunits: mTOR, RICTOR, SIN1 and mLST8. The four-residue cancer-associated 1455-EWED-1458 duplication in the FAT region of mTOR is highlighted. Also shown is a schematic sketch of the subunit arrangement within a dimeric mTORC2 complex, with the regions of interest highlighted. (**b**) Steady-state kinetic analysis of mTORC2_EWED_dup_ mutant phosphorylating full-length AKT1^ki^ (kinase-inactive mutant D274A). Inset: Zoomed in plot of mTORC2_WT to better visualise data. Reactions contained increasing concentrations of the AKT1 substrate as indicated, but an equal amount (1 µg) of total AKT1 from each reaction was loaded on a Phos-gel to achieve a linear range of detection. Gels were stained with Comassie InstantBlue stain and the intensities of the phosphorylated and nonphosphorylated bands were quantified with a ChemiDoc Touch Imaging System (Bio-Rad). Graphs show means ± SEM (with markers as indicated) of three independent experiments. Data are plotted as velocity over enzyme concentration.

**Extended Data Fig. 4:**
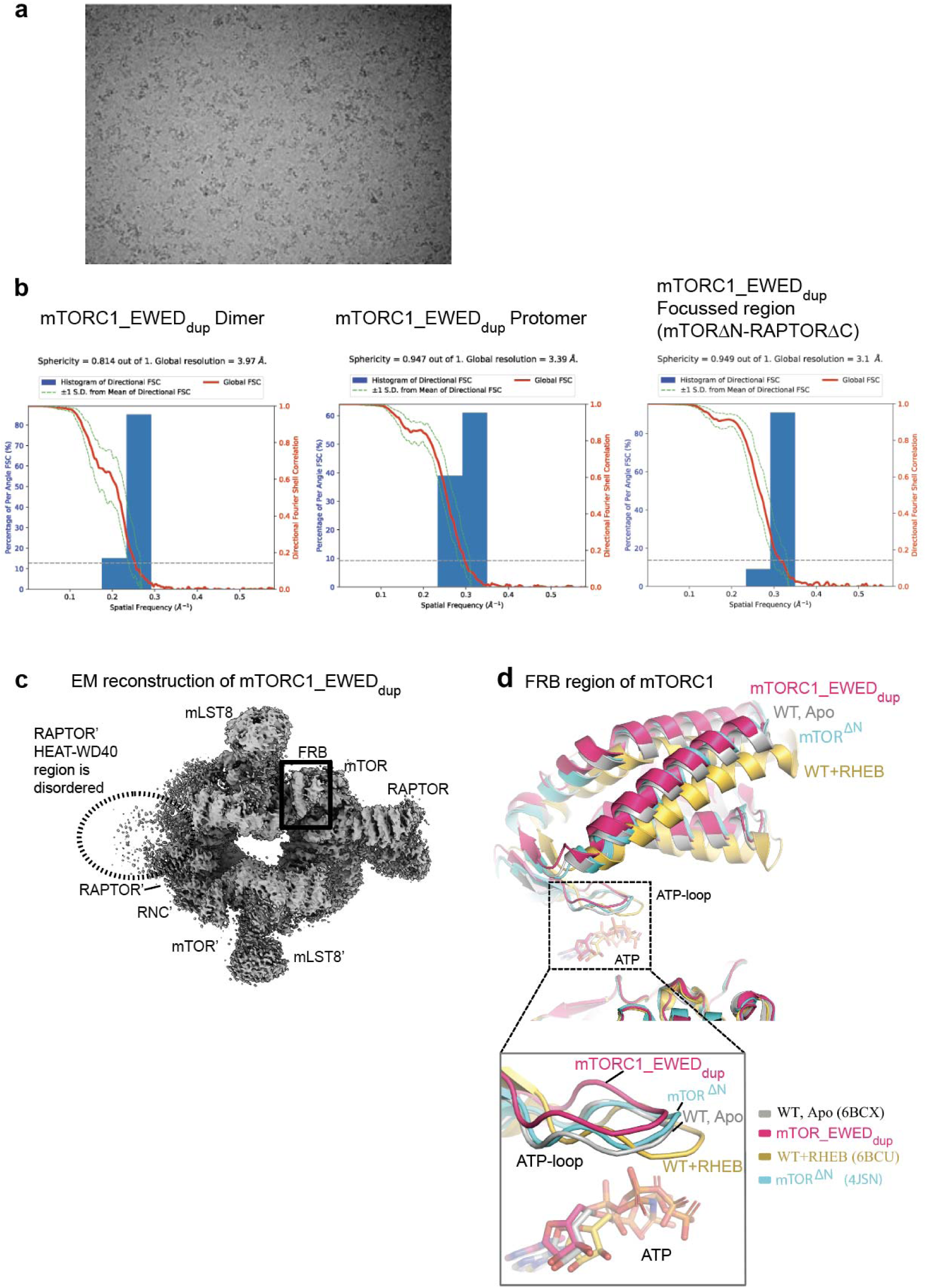
Cryo-EM structure determination of mTORC1_EWED_dup_ variant. **(a)** Top: A representative cryo-EM micrograph of mTORC1_EWED_dup_ variant. **(b)** The FSC curves for the B-factor sharpened post-processed reconstructions suggest a final resolution of the 4.0 Å for mTORC1_EWED_dup_ Dimer (sphericity 0.8), 3.4 Å for mTORC1_EWED_dup_ protomer (sphericity 0.9) and 3.1 Å for mTORC1_EWED_dup_ focused protomer region (mTORΔN-RAPTORΔC, sphericity 0.9). The global resolution and the sphericity was calculated using 3D FSC server. **(c)** Cryo-EM reconstruction of mTORC1_EWED_dup_. **(d)** Alignment of cancer-associated mTORC1_EWED_dup_ structure (dark red) with apo mTORC1_WT (6BCX, gray), RHEB-activated mTORC1_WT (6BCU, orange) and the truncated mTOR_ΔN-HEAT_-mLST8 crystal structure (4JSN, cyan). The alignment was done on the kinase C-lobe of mTOR. The FRB region from the kinase N-lobe and the close view of the ATP-loop from all structures are shown, revealing that the mTORC1_EWED_dup_ hyperactivated mutant, similarly to the partially activated mTOR_ΔN-HEAT_-mLST8 truncation variant, does not adopt the stable signature features of the activated RHEB-mTORC1, such as the realignment of the ATP-binding loop in the N-lobe with the catalytic residues in the C-lobe.

**Source data for Fig. 2b:**
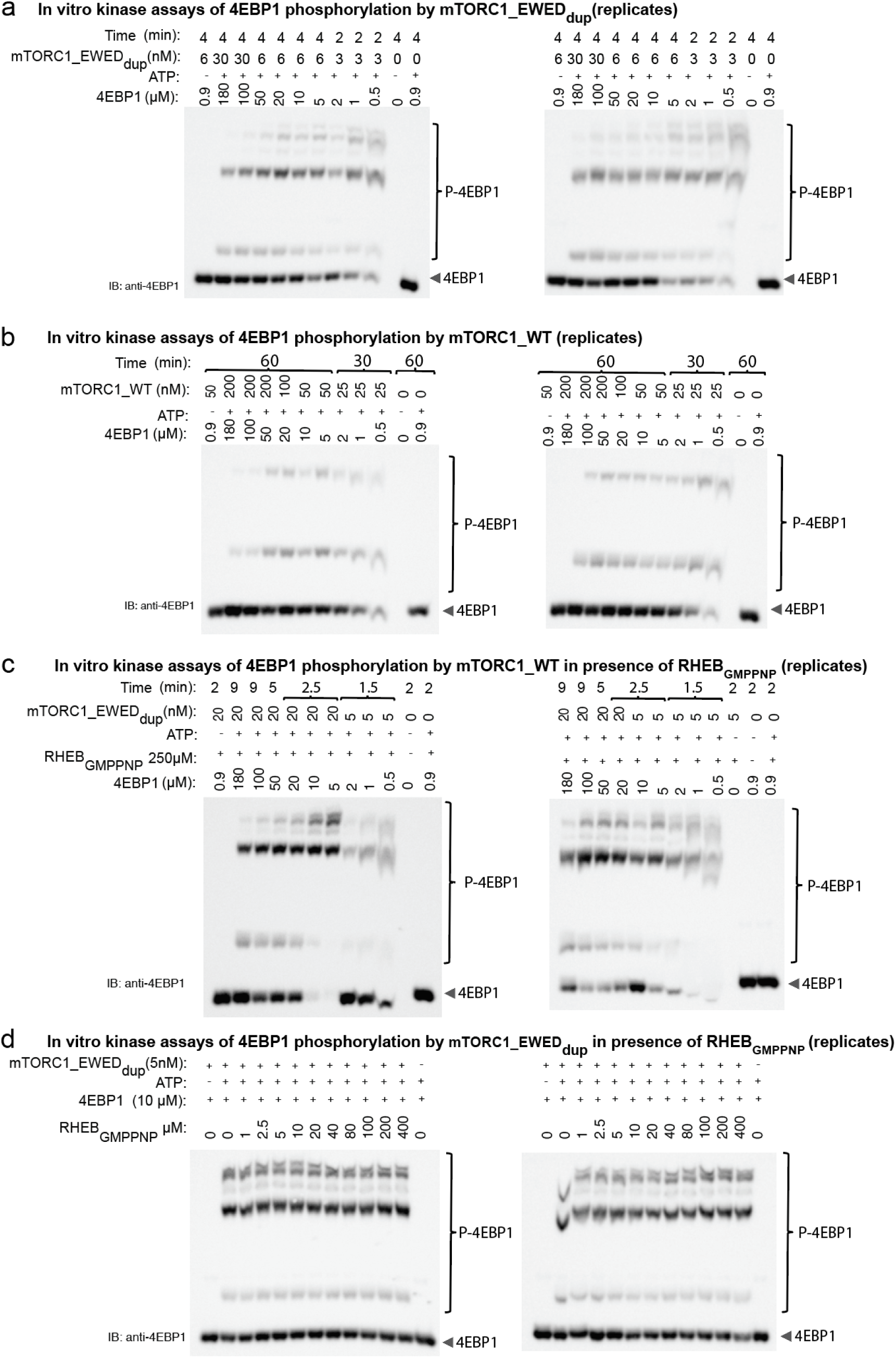
Phosphorylation of 4EBP1 by mTORC1_EWED_dup_ mutant in the absence and presence of RHEB-GMPPNP. (**a**) Immunoblots showing the phosphorylation of full-length 4EBP1 by mTORC1_EWED_dup_ mutant. Reactions contained increasing amount of the 4EBP1 substrate as indicated, but an equal amount (70 ng) of total 4EBP1 from each reaction was loaded on a Phos-gel in order to achieve a linear range of detection for quantitative Western blot analysis. Two independent experiments are shown. (**b**) Immunoblots showing the phosphorylation of 4EBP1 by mTORC1_WT. (**c**) Immunoblots showing the phosphorylation of 4EBP1 by mTORC1_WT in the presence of 250 μM RHEB-GMPPNP. (**d**) Immunoblots showing the phosphorylation of 4EBP1 by mTORC1_EWED_dup_ mutant in the presence of increasing RHEB-GMPPNP concentration as indicated. The 4EBP1 was at a constant concentration of 10 μM. RHEB does not further activate mTORC1_EWED_dup_ to phosphorylate 4EBP1. A total amount of 70 ng of total 4EBP1 was loaded on a Phos-gel. Two independent experiments are shown.

**Source data for Extended Fig. 3b.**
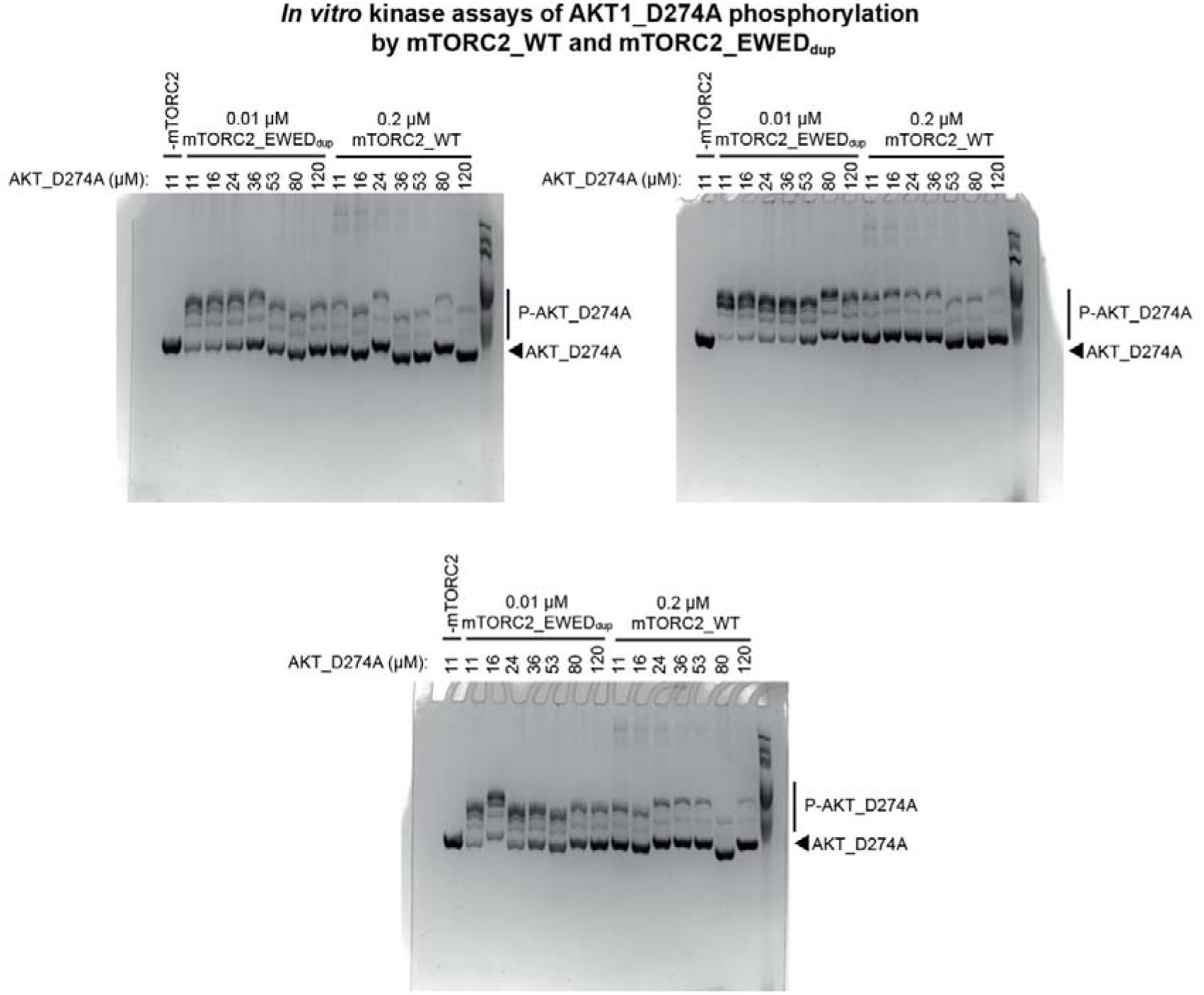
Comassie InstantBlue-stained Phos-gel showing the phosphorylation of the AKT1 substrate by mTORC2_EWED_dup_ and mTORC2_WT, as indicated. Three independent experiments are shown.

